# Genome-wide association study of Alcohol Use Disorder Identification Test (AUDIT) scores in 20,328 research participants of European ancestry

**DOI:** 10.1101/147397

**Authors:** Sandra Sanchez-Roige, Pierre Fontanillas, Sarah L. Elson, the 23 and Me Research Team, Joshua C. Gray, Harriet de Wit, Lea K. Davis, James MacKillop, Abraham A. Palmer

**Affiliations:** Department of Psychiatry, University of California San Diego, La Jolla, CA, 92093, USA; 23andMe, Inc., Mountain View, CA, USA; Department of Psychology, University of Georgia, USA; Center for Alcohol and Addiction Studies, Brown University School of Public Health, Providence, RI 02903, USA; Department of Psychiatry and Behavioral Neuroscience, University of Chicago, Chicago, IL 60637, USA; Vanderbilt Genetics Institute; Division of Genetic Medicine, Department of Medicine, Vanderbilt University, Nashville, TN, USA; Peter Boris Centre for Addictions Research, McMaster University/St. Joseph’s Healthcare Hamilton, Hamilton, ON L8N 3K7, Canada; Homewood Research Institute, Guelph, ON N1E 6K9, Canada; Institute for Genomic Medicine, University of California San Diego, La Jolla, CA, USA

## Abstract

Genetic factors contribute to the risk for developing alcohol use disorder (**AUD**). In collaboration with the genetics company 23andMe, Inc., we performed a genome-wide association (**GWAS**) study of the Alcohol Use Disorder Identification Test (**AUDIT**), an instrument designed to screen for alcohol misuse over the past year. Our final sample consisted of 20,328 research participants of European ancestry (55.3% females; mean age = 53.8, SD = 16.1) who reported ever using alcohol. Our results showed that the ‘chip-heritability’ of AUDIT score, when treated as a continuous phenotype, was 12%. No loci reached genome-wide significance. The gene *ADH1C*, which has been previously implicated in AUD, was among our most significant associations (4.4 × 10^−7^; rs141973904). We also detected a suggestive association on chromosome 1 (2.1 × 10^−7^; rs182344113) near the gene *KCNJ9*, which has been implicated in mouse models of high ethanol drinking. Using LD score regression, we identified positive genetic correlations between AUDIT score and AUD, high alcohol consumption, and cigarette smoking. We also observed an unexpected positive genetic correlation between AUDIT and educational attainment, and additional unexpected negative correlations with BMI/obesity and attention-deficit/hyperactivity disorder (**ADHD**). We conclude that conducting a genetic study using data from a population unselected for AUD and responding to an online questionnaire may represent a cost-effective strategy for elucidating the etiology of AUD.

## INTRODUCTION

The heritability of AUD and associated symptomatology such as high alcohol consumption has been estimated at ~50% by family and twin studies (Mbarek et al., 2015; Verhulst, Neale, & Kendler, 2015), with a smaller proportion being attributable to additive effects of common genetic variation (e.g. 33% for AUD (Mbarek et al., 2015); 18% (Vrieze, McGue, Miller, Hicks, & Iacono, 2013) and 13% (Clarke et al., 2017) for alcohol consumption).

The search for specific genes that convey risk for AUD has been an active area of research for several decades. There have been numerous family-based linkage studies of AUD (Agrawal & Bierut, 2012; Edenberg & Foroud, 2014; Enoch, 2013; Rietschel & Treutlein, 2013), as well as candidate gene association studies. Robust linkage signals have been found near the cluster of aldehyde dehydrogenase (*ALDH*) genes on chromosome 12 (Wall, Luczak, & Hiller-Sturmhöfel, 2016), and alcohol dehydrogenase (*ADH*) genes on chromosome 4 (Long et al., 1998; Williams et al., 1999), a result that has also been replicated by candidate gene association studies (Li, Zhao, & Gelernter, 2011, 2012; Luo et al., 2006; Macgregor et al., 2009; Thomasson et al., 1991; van Beek, Willemsen, de Moor, Hottenga, & Boomsma, 2010).

More recently, genome-wide association studies (**GWAS**) have been used to explore the genetic basis of AUD (Hart & Kranzler, 2015; Tawa, Hall, & Lohoff, 2016). The most robust and replicated risk alleles in European, African American, and Asian ancestry populations map to alcohol-metabolizing enzyme genes on chromosome 4q22-23 and 12q24: *ADH1B* (Gelernter et al., 2014; Clarke et al., 2017; Xu et al., 2015), *ADH1C* (Clarke et al., 2017; Edenberg et al., 2010; Frank et al., 2012; Gelernter et al., 2014; Treutlein et al., 2009), *ADH5* (Clarke et al., 2017), *ADH7* (Park et al., 2013) and *ALDH2* (Jorgenson et al., 2017; Park et al., 2013; Quillen et al., 2014; Takeuchi et al., 2011; Yang et al., 2013). More recent GWAS that target alcohol consumption rather than AUD have identified novel genes including *KLB*, which influence both high alcohol consumption in humans (Clarke et al., 2017; Jorgenson et al., 2017; Schumann et al., 2016) and ethanol preference in mice (Schumann et al., 2016).

In collaboration with the genetics company 23andMe, Inc., we performed a GWAS for alcohol misuse using the Alcohol Use Disorders Identification Test (**AUDIT**), a questionnaire developed to screen for alcohol misuse in the past year (Saunders, Aasland, Babor, de la Fuente, & Grant, 1993a). The estimated heritability of AUDIT score is 60%, similar to the heritability of AUD (Mbarek et al 2015). Self-reported AUDIT scores are predictive of future problematic drinking and higher AUD risk (Allen, Litten, Fertig, & Babor, 1997; Boschloo et al., 2010), perhaps because the AUDIT includes questions that are related to the criteria for AUD (e.g. DSM-V, criterion 2: “More than once wanted to cut down or <u>stop drinking</u>, or tried to, but couldn’t?” versus AUDIT, item 4: “How often during the last year have you found that you were not able to <u>stop drinking</u> once you had started?”). A previous GWAS of dichotomized AUDIT scores in 7,842 individuals in an unselected Dutch population did not reveal any significant associations (Mbarek et al 2015). Here, to maximize power and to capture the dimensionality of alcohol misuse, we treated AUDIT scores from 20,328 research participants as a continuous trait rather than dichotomizing it by using a threshold score. We hypothesized that a GWAS for AUDIT scores might identify some of the same alleles that influence AUD, even though our cohort had relatively modest AUDIT scores.

## MATERIALS AND METHODS

### Sample

All participants included in the analyses were drawn from the customer base of 23andMe, Inc., a consumer genetics company. Participants provided informed consent and answered surveys online under a protocol approved by Ethical and Independent Review Services, an independent AAHRPP-accredited institutional review board (http://www.eandireview.com). We restricted the original sample (~25,000 individuals) to a set of unrelated participants of European ancestry (>97% as determined through an analysis of local ancestry [Durand, Do, Mountain, Macpherson, 2014]; see **Supplementary** for additional details) for whom AUDIT data were available. Participants were excluded if they reported that they never drank alcohol (*N* = 1,376). The final number of participants included in the analysis was 20,328. Recruitment occurred over an approximately four-month period in 2015. Sociodemographic details are described in the **Supplementary Table 1**.

### AUDIT scores

To evaluate alcohol misuse in the past year, participants completed the AUDIT (Saunders, Aasland, Babor, de la Fuente, & Grant, 1993). We only included subjects who answered yes to the question “Have you ever in your life used alcohol” (i.e., “ever drinkers” vs “never drinkers”). The ten-item AUDIT questionnaire yields scores from 0 to 40. Since the scores were not normally distributed (by visual inspection), we used a log-10 transformation, which is frequently employed to approximate a normal distribution for AUDIT (**Supplementary Table 2**).

### Genotyping, quality control and imputation

DNA extraction and genotyping were performed on saliva samples by the National Genetics Institute, a CLIA-certified laboratory. Samples were genotyped on 23andMe custom genotyping array platforms (Illumina HumanHap550+ Bead chip V1 V2, OmniExpress+ Bead chip V3, Custom array V4). Quality control of genetic variants and imputation were performed by 23andMe (see **Supplementary Table 3**). A full description of the methods have been reported elsewhere (Hyde et al., 2016; Lo et al., 2016).

### Estimation of variance in AUDIT scores explained by the genotyped SNPs

To estimate the proportion of phenotypic variance explained (‘chip heritability’; h_g_^2^), we used a genomic restricted maximum likelihood (GREML) method implemented in Genetic Complex Trait Analysis (GCTA; Yang, Lee, Goddard, & Visscher, 2013). In brief, the GREML method estimates the proportion of variation in a phenotype that is due to all SNPs, and exploits the fact that genotypic similarity (i.e., “relatedness”, measured using genotyped SNPs) will be correlated with phenotypic similarity for heritable traits. Individual-level quality control was implemented, and distantly related individuals with pair-wise relationships were filtered at two thresholds (K_IBS_ < 0.05 and K_IBS_ < 0.03). We included age (inverse-normalized), self-reported sex (male/female), genotyping platform and top four principal components as covariates. GREML analyses were run using only directly genotyped SNPs to construct the GRM.

### Chip-heritability using LD Score Regression

We used a second method to measure chip heritability of AUDIT that is implemented by Linkage Disequilibrium Score Regression Coefficient (LDSC; Bulik-Sullivan et al., 2015a). To standardize the input file (GWAS summary statistics), we followed quality controls as implemented by the LDSC python software package. We used pre-calculated LD scores (“eur_w_ld_chr/” files (Finucane et al., 2015); MHC region excluded) for each SNP using individuals of European ancestry from the 1000 Genomes project, suitable for LD score analysis in European populations. We restricted the analysis to well-imputed SNPs: the SNPs were filtered to HapMap3 SNPs (International HapMap 3 Consortium et al., 2010), and were required to have a minor allele frequency (**MAF**) above 1%. InDels, structural variants, strand-ambiguous SNPs, and SNPs with extremely large effect sizes (χ^2^ > 80) were removed. In addition, this approach allowed us to distinguish between genomic inflation attributed to polygenic signal, from confounding biases such as population stratification or polygenicity (LD Score regression intercept > 1; Bulik-Sullivan et al., 2015a; Bulik-Sullivan et al., 2015b). As expected under polygenicity, we observed inflation of the median test statistic (Mean χ^2^ = 1.05), and adjusted for a genomic control inflation factor λ (the ratio of the observed median χ^2^ to that expected by chance) = 1.02. LD score intercept of 1.01 (SE = 0.01) suggested that deviation from the null was due to a polygenic structure rather than inflation due to population structure biases.

### Genome-wide association analysis

For quality control of genotyped GWAS results, we removed SNPs with MAF of < 0.1%, a Hardy-Weinberg *P* <10^−20^ in Europeans, or a call rate of < 90%. We also removed SNPs that were only genotyped on the 23andMe V1 platform, due to limited sample size, and SNPs on chrM or chrY. Using trio data, we removed SNPs that failed a test for parent-offspring transmission; specifically, we regressed the child’s allele count against the mean parental allele count and removed SNPs with fitted β < 0.6 and *P* < 10^−20^ for a test of β < 1. We also tested genotyped SNPs for genotype date effects, and removed SNPs with P < 10^−50^ by ANOVA of SNP genotypes against a factor dividing genotyping date into 20 roughly equal-sized buckets. For imputed GWAS results, we removed SNPs with average r^2^ < 0.50 or minimum r^2^ < 0.30 in any imputation batch, as well as SNPs that had strong evidence of an imputation batch effect. The batch effect test is an F test from an ANOVA of the SNP dosages against a factor representing imputation batch; we removed results with P < 1 × 10^−50^. We also removed linear regression results for SNPs with MAF < 0.1% because tests of low frequency variants can be sensitive to violations of the regression assumption of normally distributed residuals. We performed association tests by linear regression assuming an additive model. We included age (inverse-normal transformed), sex, the top four principal components of genotype, and indicator variables for genotype platforms as covariates (**Supplementary Table 4**).

### Phenotypic and genetic correlation analyses

We examined two distinct types of correlations: phenotypic correlations, where both variables were measured in the same individuals, and genetic correlations, where we used AUDIT data from this cohort in conjunction with summary statistics for GWAS conducted in other cohorts (**Supplementary Table 9**). The interpretation of these is different, since phenotypic correlations can be due to a combination of genetic and non-genetic factors, whereas genetic correlations measure only genetically driven correlations.

We used bivariate correlations to examine the direct phenotypic correlations between AUDIT and several variables of interest (age, gender, race, education, annual household), and to identify significant covariates for inclusion in GWAS analysis (**Supplementary Table 5**).

We calculated genetic correlations (*r*_g_) between AUDIT and 30 other complex traits or diseases using LDSC. References for the datasets used are identified in **Supplementary Table 9**. Files were standardized using the steps described in the section above (“Chip-heritability using LD Score Regression”). We did not constrain the intercepts in our analysis because the degree of sample overlap was unknown. We used False Discovery Rate (FDR) to correct for multiple testing (Benjamini & Hochberg, 1995).

### Query for expression quantitative trait loci (eQTL)

We queried eQTL evidence for our top (P < 10^−7^) GWAS SNPs using public online resources. We used the Genotype-Tissue Expression Portal (GTEx) to identify eQTLs associated with the SNPs; and the RegulomeDB (Boyle et al., 2012) to identify regulatory DNA elements in non-coding and intergenic regions of the genome in normal cell lines and tissues.

## RESULTS

### Demographics

Demographic data are shown in **Supplementary Table 1**. Mean age was 53.8 years (SD = 16.1), and 55.3% were women. The annual household income ranged from less than $14,999 (13.5%) to greater than $75,000 (21.5%), and the mean years of education completed was 16.8 (SD = 2.6). About half of the participants (49.3%) were married/partnered. Participants showed low to moderate alcohol use, average frequency of alcohol use was ~9 days per month (mean = 8.78, SD = 9.82); during the period of heaviest lifetime use, subjects reported reaching an average of 13.78±10.96 days over a 30-day period. Over the prior year, 78% of the participants reported drinking 1 or 2 drinks on a single day, and only 28% reported drinking more than 6 drinks on one occasion. Also over the prior year, 92% of the participants were able to stop drinking once they started, and 85% drank alcohol without feeling guilt or remorse.

### AUDIT scores

The distribution of the AUDIT scores is shown in **Supplementary Table 2**. The phenotypic correlations between AUDIT and demographic variables measured in the same cohort are shown in the **Supplementary Table 4**. Age and sex were negatively correlated with AUDIT scores; younger individuals and males showed higher AUDIT scores (*r* = −0.15, *P* < 0.0001; *r* = −0.17, *P* < 0.0001, respectively). BMI was *negatively* associated with AUDIT (*r* = −0.07, *P* < 0.0001), whereas household income was positively correlated with AUDIT (*r* = 0.07, *P* < 0.0001). AUDIT scores were slightly higher in unmarried individuals (*r* = −0.02, *P* = 0.013) but we did not observe significant correlations with years of education (*r* = 0.01, *P* = 0.085). Measures of alcohol use were positively correlated with AUDIT scores (*r* = 0.50-0.52, *P* < 0.0001).

### Chip-heritability estimates

We estimated the chip-heritability of AUDIT at 12.05% (± 1.91%, *P* = 2.70 x 10^−11^), which is lower than previous chip-heritability estimates based on dichotomized AUDIT data (30% ±12%; Mbarek et al., 2015), and considerably lower than twin based heritabilities of alcohol abuse, dependence and alcoholism (~50%, (Enoch, 2013; Goldman, Oroszi, & Ducci, 2005).

### GWAS of AUDIT

The Manhattan and quantile-quantile (**QQ**) plots for AUDIT are shown in **Figure 1** and **Supplementary Figure 4**, respectively. The most significant association was at rs182344113, located on chromosome 1 (*P* = 2.10 × 10^−7^; β = 0.168, SE = 0.03; MAF = 0.002; **Supplementary Fig. 1**). The association was in an intergenic region of the gene *PIGM*, and near *KCNJ9* (*GIRK3*), which has been implicated in preclinical models of ethanol sensitivity. G-protein-gated inwardly rectifying potassium (GIRK) channels, which are coupled to GABA-B receptors, can be activated by ethanol (Aryal, Dvir, Choe, & Slesinger, 2009; Bodhinathan & Slesinger, 2013). Interestingly, *Kcnj9* knock-out mice exhibit excessive alcohol drinking (Dere et al., 2015).

**Figure 1.**
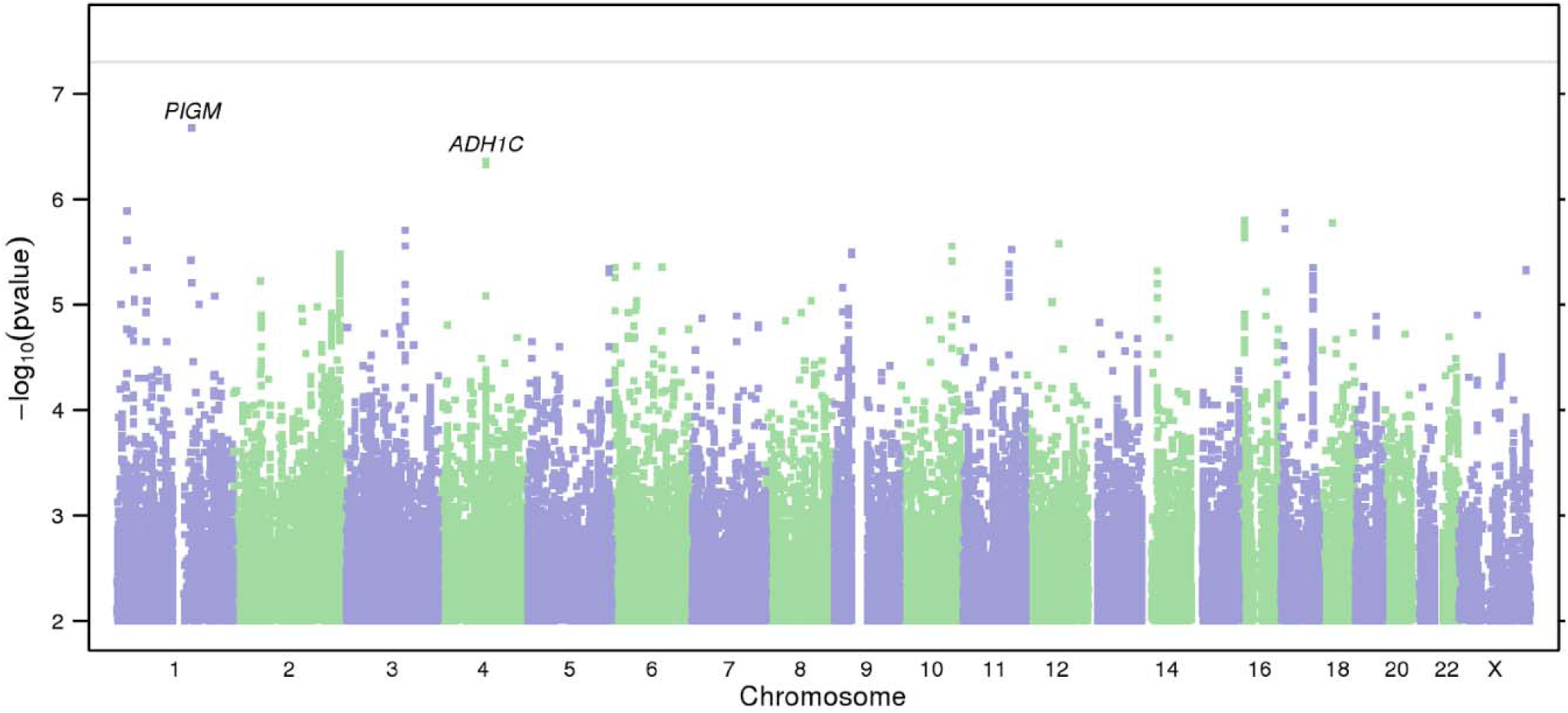
Results of GWAS on AUDIT. (**a**) Manhattan plot of GWAS results indicating the strongest associations between the 22 autosomes, X chromosome, and AUDIT. Line denotes genome-wide significance (*P* < 5 × 10^−8^).

Several other SNPs also showed suggestive associations (**Supplementary Table 6**), including rs141973904 (*P* = 4.40 × 10^−7^, β = −0.05, SE = 0.01; MAF = 0.02; **Supplementary Fig. 2**) in an intron of *ADH1C*, replicating previous findings for that same SNP in a GWAS of alcohol consumption in males (Clarke et al., 2017), and broadly consistent with numerous previous genetic studies of AUD (Biernacka et al., 2013; Edenberg, 2007; Hn et al., 2013; Thomasson et al., 1991).

Another suggestive association was at rs8059260 (*P* = 1.6 × 10^−6^, β = 0.017, SE = 0.004; MAF = 0.160; **Supplementary Fig. 3**), which is near the first exon of *CLEC16A*. Using the Genotype-Tissue Expression Portal (**GTEx**) database, we identified a *cis-* expression quantitative trait loci (**eQTLs**) for *CLEC16A* that co-localized with rs8059260 (r^2^ > 0.79; see **Supplementary Table 7**). We also found evidence of regulatory elements associated with rs8059260 using the RegulomeDB (Boyle et al., 2012; **Supplementary Table 7**).

### Previously studied candidate genes

Our results did not strongly support any of the previously published candidate gene studies of AUD (reviewed in Bühler et al., 2015; **Supplementary Table 8**); various differences including the distinction between AUD and AUDIT, demographics characteristics and especially the low prevalence of AUD in our cohort could partially account for the lack of replication.

### Genetic correlations

LD score regression (Bulik-Sullivan et al., 2015b) showed a genetic overlap between AUDIT and numerous traits (**Fig. 2** and **Supplementary Table 9**). Both alcohol consumption and AUD showed the steepest correlations with AUDIT score (r_g_ = 0.68 for both), while the better powered alcohol consumption trait (N = 70,460) yielded a significant result (*P* = 3.40 × 10^−3^) the *P* value for the genetic correlation with AUD (N = 7,280) fell just short of nominal significance (*P* = 6.41 × 10^−2^). We detected a significant negative genetic correlation between AUDIT and ADHD (r_g_ = −0.29; *P* = 1.43 × 10^−3^). We observed a positive genetic correlation between AUDIT and lifetime tobacco use (r_g_ = 0.42; *P* = 1.52 × 10^−3^). Unexpectedly, we identified a *positive* correlation between AUDIT and years of education (r_g_ = 0.27; *P* = 3.14 × 10^−5^), college attainment (r_g_ = 0.26; *P* = 8.11 × 10^−3)^ and childhood IQ (r_g_ = 0.42; *P* = 6.26 × 10^−3^). Also surprisingly, AUDIT was *negatively* genetically correlated with BMI (r_g_ = −0.25; *P* = 1.48 × 10^−4^) and adulthood obesity (r_g_ = −0.23; *P* = 2.06 × 10^−3^). Height, which is not strongly influenced by individual behavior and thus can be viewed as a negative control, was not genetically correlated with AUDIT (r_g_ = 0.02; *P* = 7.12 × 10^−1^).

**Figure 2.**
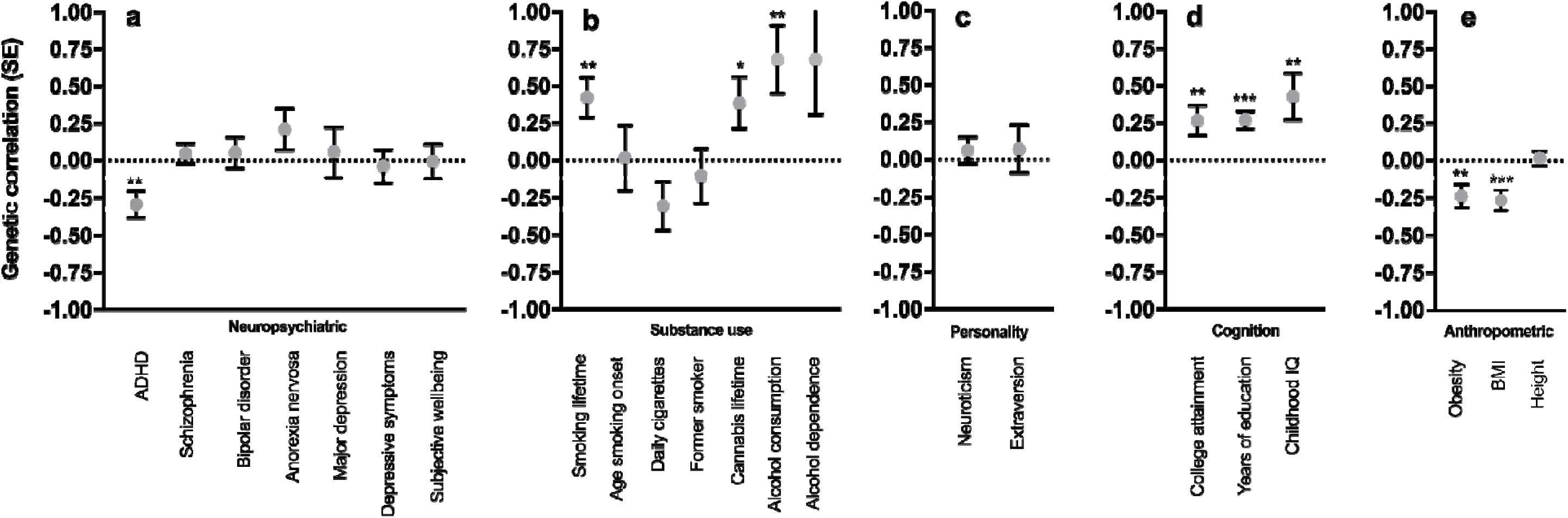
Genetic correlations between AUDIT and several traits: (**a**) neuropsychiatric, (**b**) smoking, (**c**) personality, (**d**) cognition, (**e**) anthropomorphic. * P < 0.05, ** P < 0.01, *** P < 0.0001.

## DISCUSSION

With over 20,000 research participants, ours is by far the largest genetic study of AUDIT. By using a self-report measure of alcohol misuse, as opposed to recruiting a clinically-diagnosed population, we were able to rapidly and inexpensively ascertain a large number of participants. We identified rs141973904 (**Supplementary Fig. 2**) in the *ADH* cluster on chromosome 4q23, which has been previously associated with AUD (Edenberg et al., 2010; Frank et al., 2012a; Gelernter et al., 2014; Treutlein et al., 2009a). The same SNP has recently been associated (*P* = 1.22 × 10^−15^) with alcohol consumption using 53,089 males of European ancestry (Clarke et al., 2017). Furthermore, the most associated signal, rs182344113, which resides near the *KCNJ9* (*GIRK3*) gene, was unknown, and is consistent with mouse studies of the homologous gene. The signal at rs182344113 was not significant and it will have to be replicated. We also identified a number of genetic correlations that have behavioral precedents, such as lifetime tobacco use, and several others that were unexpected, including lower BMI and obesity rates, and higher education. We found that AUDIT was genetically correlated with AUD and alcohol consumption, suggesting that a non-clinical population can be used as an alternative approach to study the genetics of AUD.

We identified modestly associated variants in alcohol metabolizing genes. Genes influencing pharmacokinetics have previously been identified through linkage, candidate gene and genome-wide association studies for AUD and related traits (reviewed in Tawa et al., 2016). The most robust signal was located in the *ADH1C* gene, which contributes to ethanol oxidation. This signal was also identified in earlier GWAS studies for alcohol consumption (Clarke et al., 2017) and AUD status in both European (Clarke et al., 2017; Frank et al., 2012) and African American (Gelernter et al., 2014) populations, suggesting that pharmacokinetic factors are an important contributor to differences in both AUDIT score and AUD.

In addition to SNPs in the alcohol metabolizing genes, linkage and candidate gene studies have identified the *GABRA2, OPRM1*, *DRD2* and *ANNK1* genes, as candidate genes associated with AUD phenotypes (Bühler et al., 2015). However, we did not find robust signals for any of them (**Supplementary Table 8**), suggesting that previous studies may have overestimated the effects of these genes, or that these genes are associated with AUD but not AUDIT.

The strongest association we observed resides near *KCNJ9*; the frequency of the implicated allele was very low (MAF = 0.002). *KCNJ9* encodes one of the G protein-activated inwardly rectifying K^+^ channels (GIRK3), which are expressed in the brain (Aguado et al., 2008; Koyrakh et al., 2005), and can be directly activated by ethanol (Herman et al., 2015), even at low concentrations. In humans, two linkage studies have mapped this region for AUD (Hill et al., 2004), age of onset of drinking, harm avoidance, and novelty seeking (Dick et al., 2002). Additionally, DNA methylation levels of CpG in the promoter region of the *GRIK3* gene showed altered expression in postmortem prefrontal cortex tissue of male alcoholics (Wang, Xu, Zhao, Gelernter, & Zhang, 2016). In mice, *Kcnj9* also harbors a QTL for a variety of alcohol-related behaviors, including: ethanol preference (Tarantino, McClearn, Rodriguez, & Plomin, 1998), ethanol aversion (Risinger & Cunningham, 1998), acute sensitivity to ethanol (Tipps, Raybuck, Kozell, Lattal, & Buck, 2016), and hypersensitivity to ethanol withdrawal (Kozell, Walter, Milner, Wickman, & Buck, 2009). Mice lacking GIRK3 in the brain have elevated alcohol drinking, without affecting the sensitivity to ethanol intoxication (Tipps, Raybuck, Kozell, Lattal, & Buck, 2016). Collectively, these results could provide an example of convergent results from humans and mice; however, until this non-significant observation is replicated it should be viewed with caution.

We hypothesized that the genetic risk for AUD is likely to overlap with numerous traits relevant to addiction and psychiatric phenotypes, based on previous epidemiological data (Compton, Thomas, Stinson, & Grant, 2007), twin studies (Kendler, Heath, Neale, Kessler, & Eaves, 1993; Knopik, Heath, Bucholz, Madden, & Waldron, 2009; Pickens, Svikis, McGue, & LaBuda, 1995) and recent genetic correlations between alcohol consumption and neuropsychiatric traits (Clarke et al., 2017). We showed positive genetic correlations between AUDIT and lifetime cigarette smoking, as previously observed between alcohol consumption and daily cigarettes, and tobacco initiation (Nivard et al., 2016; Vink et al., 2014). We also observed shared genetic architecture across AUDIT and two other AUD traits: alcohol consumption (Schumann et al., 2016), and AUD diagnosis (*P* = 0.062; Gelernter et al., 2014). With regards to other psychiatric traits, we found a negative correlation with ADHD; but there were no other significant genetic correlations between AUDIT and psychiatric traits. This result is generally consistent with a recent GWAS by Clarke and colleagues (2017), where they did not observe strong correlations between alcohol consumption and psychiatric conditions, with the exception of schizophrenia. This difference between our results and those of Clarke et al (2017) may be due to differences in cohorts used.

Unexpectedly, we found *positive* genetic correlations between AUDIT and years of education, college attainment and childhood IQ. This association was suggestive for the within-sample phenotypic correlation (**Supplementary Table 5**) and was significant for the genetic correlation (**Supplementary Table 9**). Consistent with this finding, Clarke and colleagues (2017) reported that college attainment and years of education were positively genetically correlated with alcohol consumption in females but not males.

Also unexpectedly, we observed *negative genetic* correlations between AUDIT and BMI and obesity. We also observed a *phenotypic* correlation between high AUDIT scores and low BMI. Previous studies have shown both positive and negative phenotypic correlations between alcohol use and BMI and obesity (Breslow & Smothers, 2005; Green et al., 2016; Hn et al., 2013; Sobczyk-Kopciol et al., 2011; Tolstrup et al., 2005), which may reflect differences in the populations used. The generalizability and biological meaning of these observations will require further research.

Our study is not without limitations. Cumulative AUDIT scores reflect two distinct constructs: one measuring alcohol consumption and another measuring alcohol-related problems; thus, AUDIT scores may conflate multiple genetic signals (Bergman & Källmén, 2002; Shevlin & Smith, 2007). In addition, our study focused on a cohort with relatively low levels of alcohol use; the unexpected positive genetic correlation between AUDIT and educational attainment, and the negative genetic correlation between AUDIT and both BMI/obesity and ADHD, may not generalize to cohorts with higher levels of alcohol use (Goldman et al., 2005). Another limitation of this study is the reliance on self-reported alcohol consumption, which may have induced biases and result in a more heterogeneous sample (Agrawal et al., 2012). Finally, AUDIT explicitly asks about alcohol use in the past year (i.e. state rather than trait), this temporal specificity is suboptimal for a genetic study.

Although we used a unique and potentially powerful technique to examine the genetic basis of alcohol misuse, we recognize that alcohol consumption, misuse and dependence are influenced by numerous factors, both genetic and environmental (e.g. availability of alcohol, social norms, laws, psychosocial and personality factors, expectancies, health factors). Further, individuals vary in susceptibility at every stage of alcohol use from initiation to severe dependence, including the continued use after a first drink, the direct subjective and behavioral effects of the drug, withdrawal severity, tolerance and susceptibility to relapse, among others. Genetic factors are likely to influence variability at each of these stages, but different stages may be influenced by different sources of genetic variability. Thus, whereas we have investigated genetic variance related to an intermediate outcome measure of alcohol misuse (as measured by the AUDIT), it remains to be determined exactly how genetic sources of variation influence alcohol consumption. This ambiguity limits our ability to elucidate the underlying molecular mechanisms identified by GWAS of alcohol use and abuse.

Nonetheless, unlike studies of disease traits, which require careful diagnosis and ascertainment, we rapidly obtained a large cohort for which genotype data were available. We replicated a previously identified signal (*ADH1C*), and identified a novel GWAS signal (near *KCNJ9*) that has preclinical correlates. Our approach shows that genetic studies of AUDIT in community-based samples are an economical and effective alternative to rigorously diagnosed AUD cohorts that can nevertheless be used to gain insight into the biology of AUD, and comorbid psychopathology.

## ACKNOWLEDGEMENTS

We would like to thank the 23andMe research participants and employees for making this work possible.

Supplementary Information with nine additional tables are available on the *Neuropsychopharmacology* website

---

**Supplementary Figure 1.**
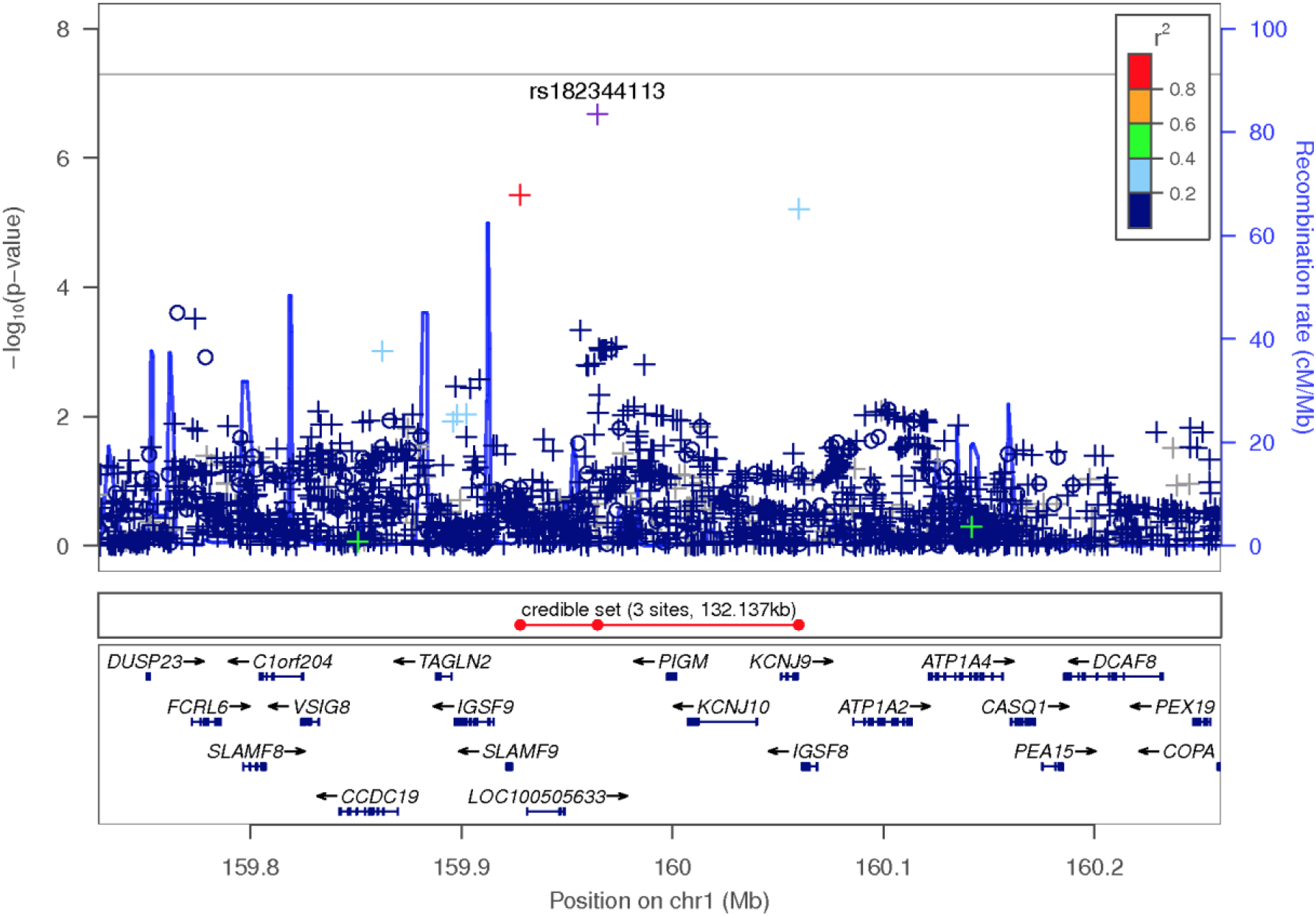
Regional association plot focusing on top SNP rs182344113 on chromosome 1.

**Supplementary Figure 2.**
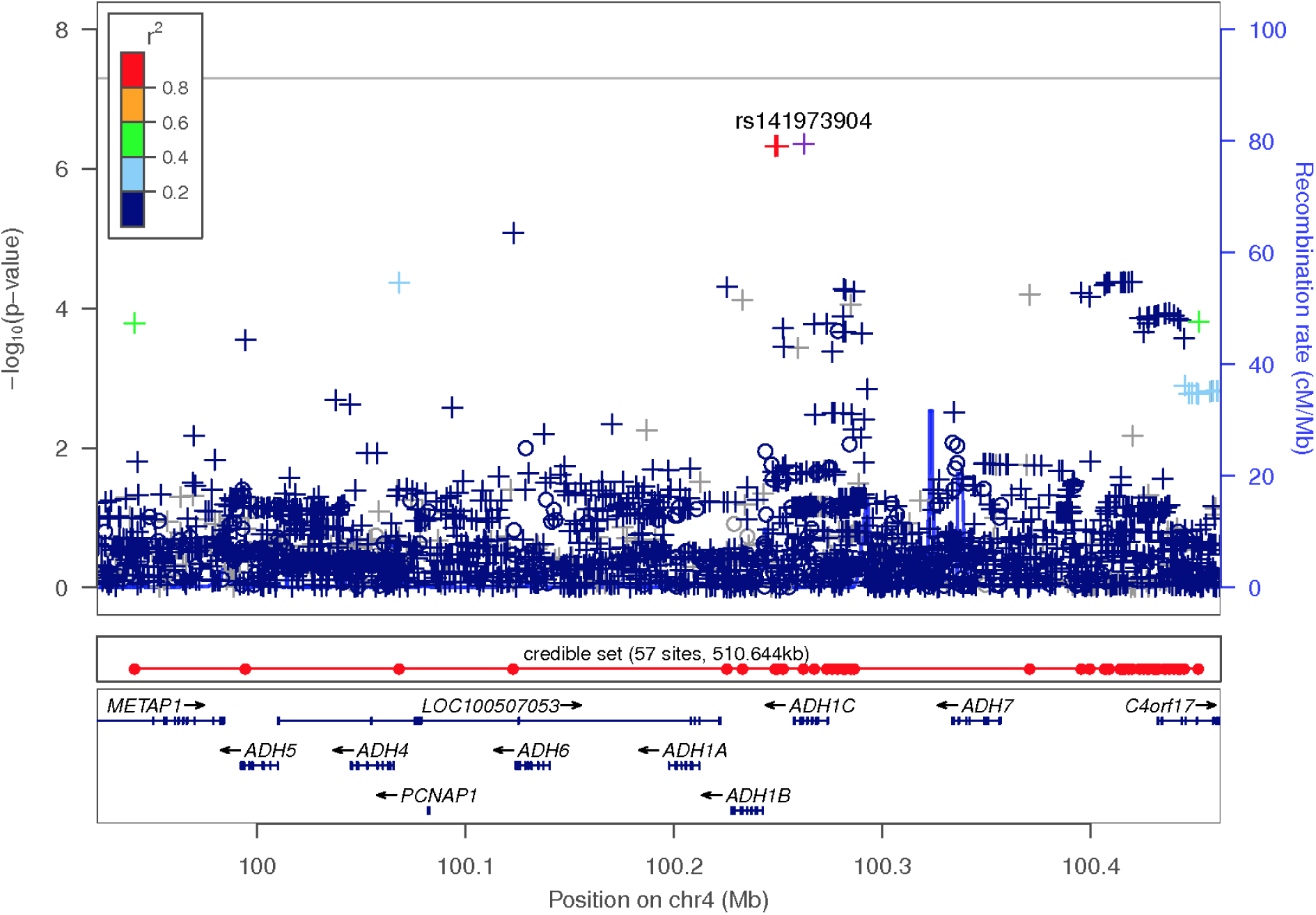
Regional association plot showing the second index SNP, rs141973904, located in the gene *ADH1C* on chromosome 4.

**Supplementary Figure 3.**
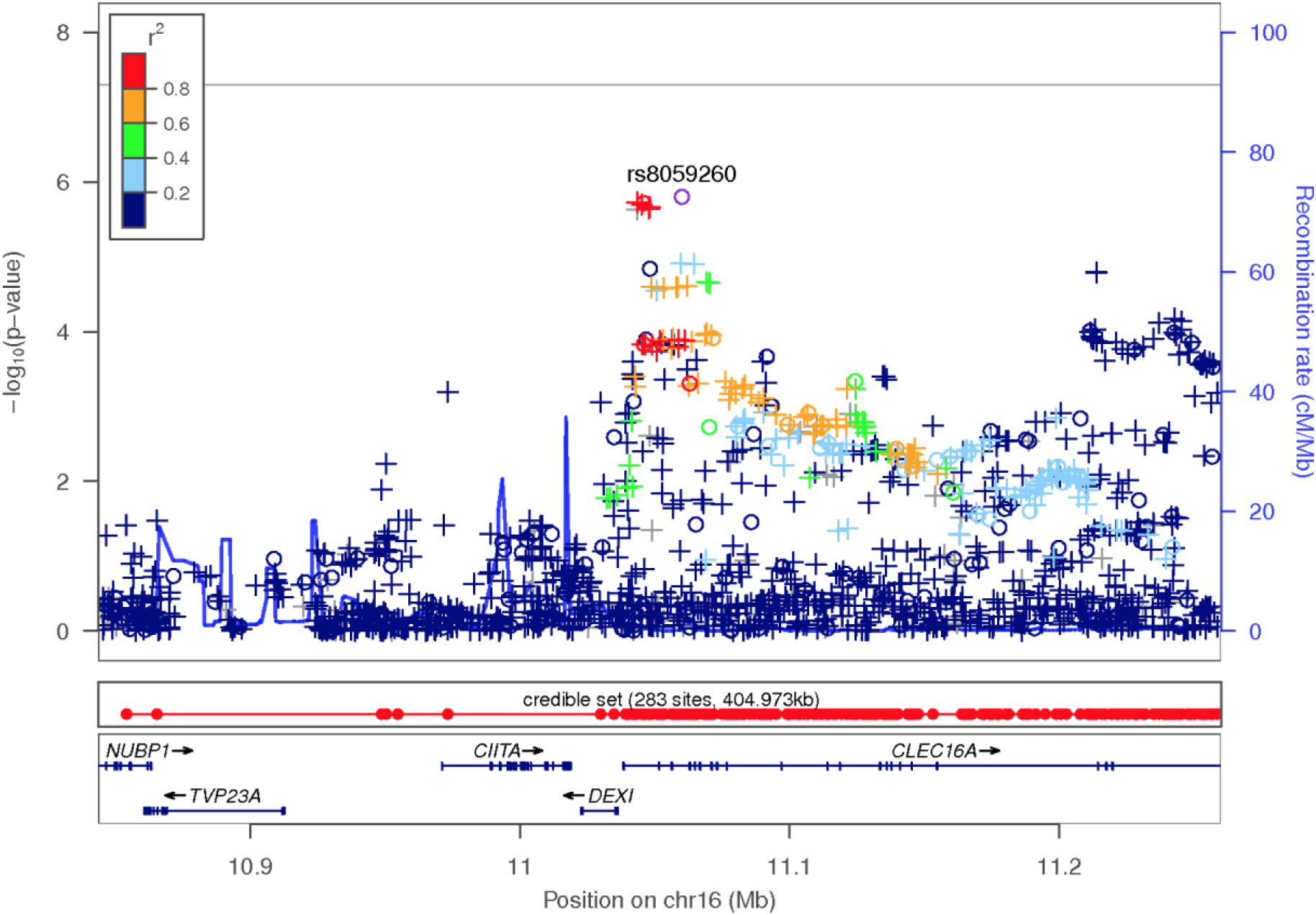
Regional association plot showing the index SNP rs8059260, located in the gene *CLEC16A* on chromosome 16.

**Supplementary Figure 4.**
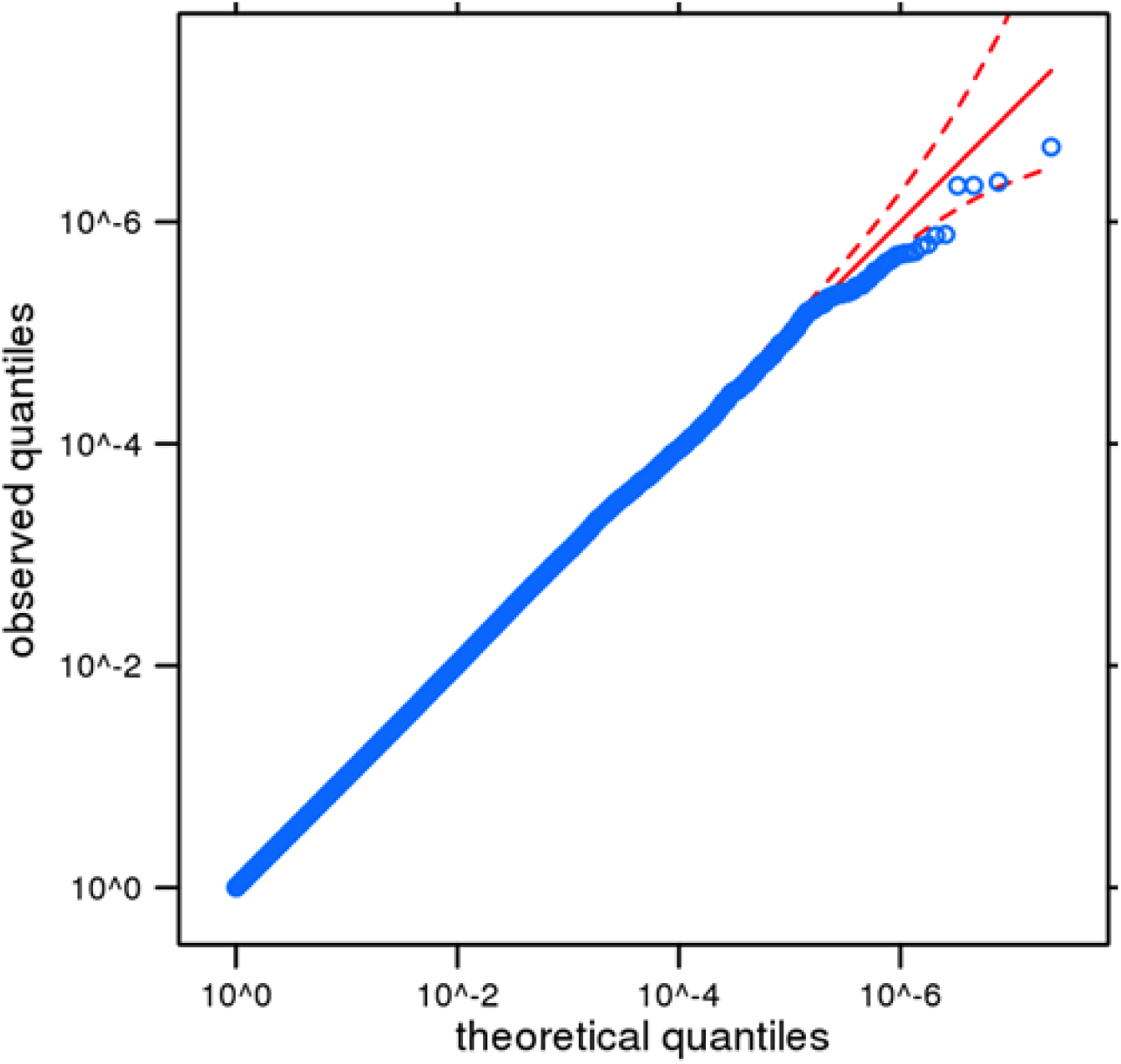
Quantile-Quantile (QQ) plot of AUDIT. The results have been adjusted for a genomic control inflation factor λ=1.021 (sample size = 20,328).

**Supplementary Table 1.** Demographic characteristics of the 23andMe cohort

|  | 23andMe Cohort * | N ** |
| --- | --- | --- |
| **Demographics** |  |  |
| Age in years (mean, S.D) | 53.79 (16.08) | 23,677 |
| Age in years (range) | 18-101 | 23,677 |
| Female | 55.30% | 23,677 |
| BMI (mean, S.D) | 27.00 (5.79) | 22,889 |
| Years of education (mean, S.D) | 16.75 (2.62) | 20,460 |
| Household income (mean, S.D) | 6.07 (1.98) | 17,919 |
| Marital status (married vs. unmarried) | 49.30% | 21,862 |
| **Alcohol use** <sup>1</sup> |  |  |
| Alcohol lifetime use (N, never/ever) | 1,376 / 23,108 | 24,484 |
| Days of alcohol use (past 30 days) | 8.78 (9.82) | 21,727 |
| Days of alcohol use (heaviest, lifetime, 30-day period) | 13.78 (10.96) | 21,229 |

*Abbreviations* \* European Ancestry only; \*\* N prior to GWAS analysis (final sample size = 20,328); N sample size; *BMI* Body Mass Index; *Household income* 9 categories from 10K to 500K

**Supplementary Table 2.**
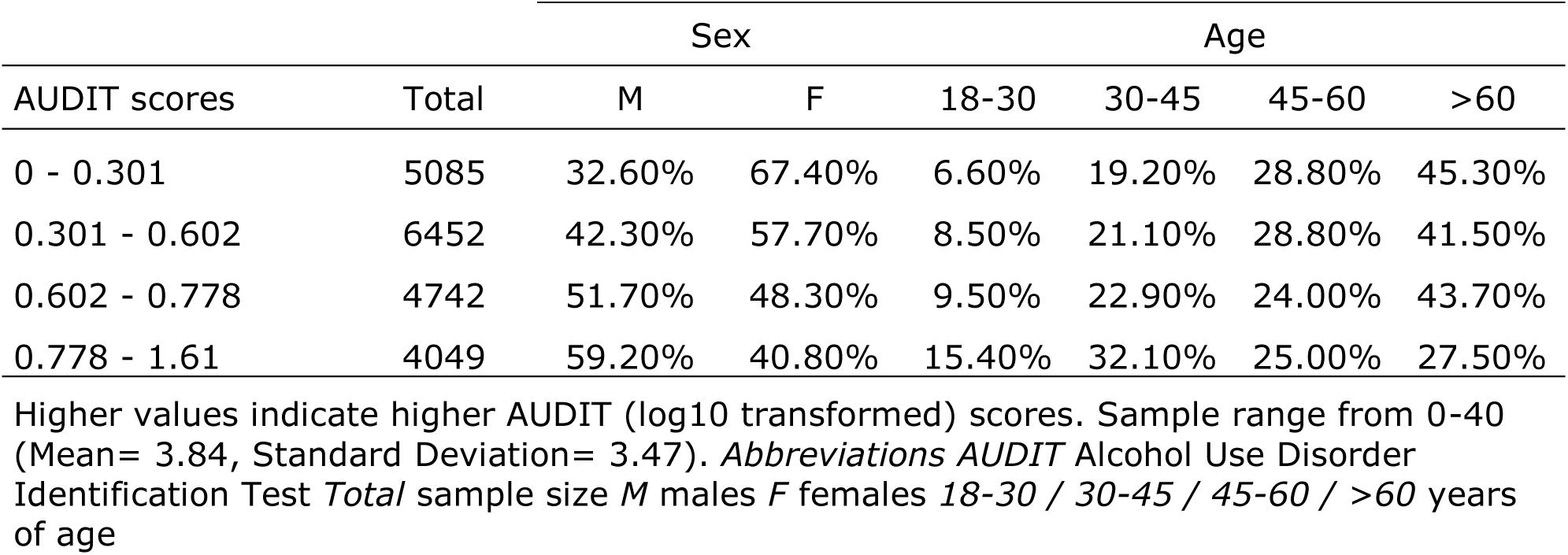
Distribution (%) of AUDIT scores

**Supplementary Table 3.** Quality Control filters at SNP-level for genotyped and imputed dosage data, and association test results data

|  | Genotyped | Imputed Dosage | Association Results |
| --- | --- | --- | --- |
| no filters | 1,050,074 | 15,574,220 | - |
| not V1/V2-only platforms, chrM, chrY | 1,015,755 | - | - |
| parent-offspring test | 1,012,649 | - | - |
| Minor Allele Frequency > 0% | 1,008,228 | 15,574,220 | - |
| Hardy-Weinberg Equilibrium > 1e-20 | 954,373 | - | - |
| genotype call rate > 90% | 934,855 | - | - |
| imputation quality | - | 13,230,301 | - |
| batch effects | 926,512 | 13,204,909 | - |
| imputed only | - | - | 12,354,282 |
| both passed | - | - | 13,204,463 |
| genotyped only | - | - | 13,280,794 |
| no test result | - | - | 12,010,653 |
| Minor Allele Frequency < 0.1% * | - | - | 11,508,740 |

*Notes \* poor residual behavior*

**Supplementary Table 4.**
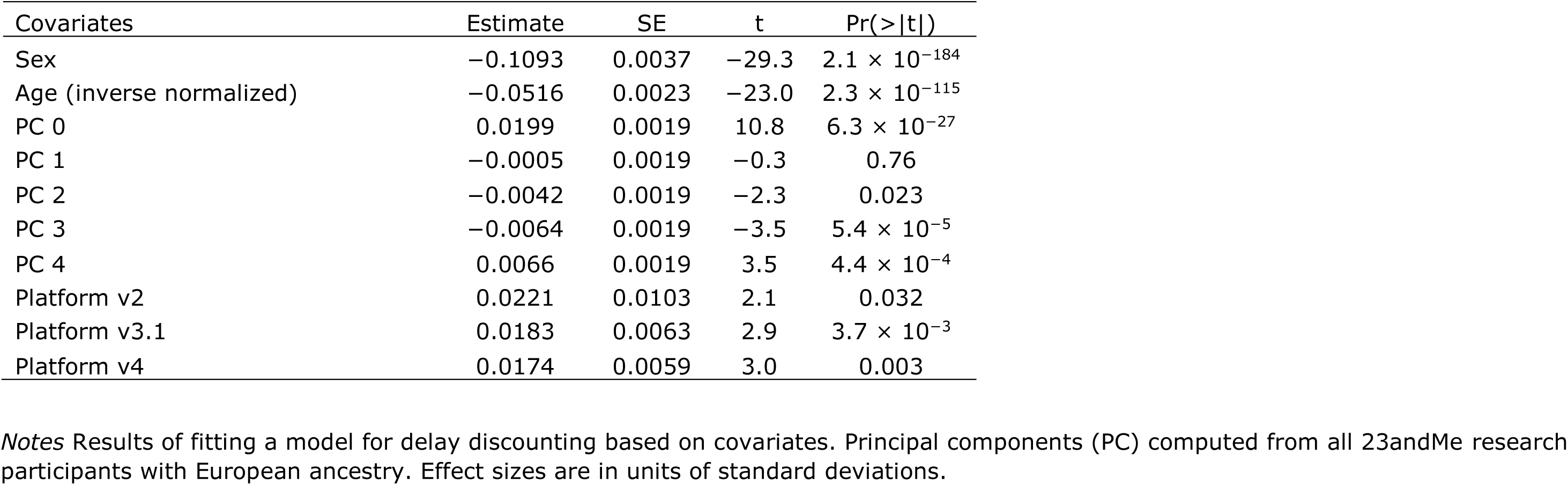
Null model with covariates

**Supplementary Table 5.** Phenotypic correlations between AUDIT scores and demographic, drug (alcohol, tobacco, marijuana; current/heaviest use) and caffeine use, and impulsivity traits

| Phenotype | Pearson *r* | *P* value | N |
| --- | --- | --- | --- |
| **Demographics** |  |  |  |
| Age in years * | -0.15 | $4.71 \times 10^{-109}$ | 20,746 |
| Sex * | -0.17 | $3.83 \times 10^{-136}$ | 20,754 |
| BMI * | -0.07 | $1.46 \times 10^{-21}$ | 20,098 |
| Household income * | 0.07 | < 0.0001 | 15,818 |
| Marital status | -0.02 | 0.013 | 19,177 |
| Years of education | 0.01 | 0.085 | 17,948 |
| **Alcohol Use** |  |  |  |
| Days of use (Past 30 days) * | 0.52 | < 0.0001 | 20,185 |
| Days of heaviest use (Lifetime, 30-days period) * | 0.50 | < 0.0001 | 20,521 |

Notes Pairwise phenotypic correlations (*r*) Abbreviations \* *P* < 0.001 \*\* Prior to GWAS analysis (Final sample size (*N*) = 20,328)

**Supplementary Table 6.**
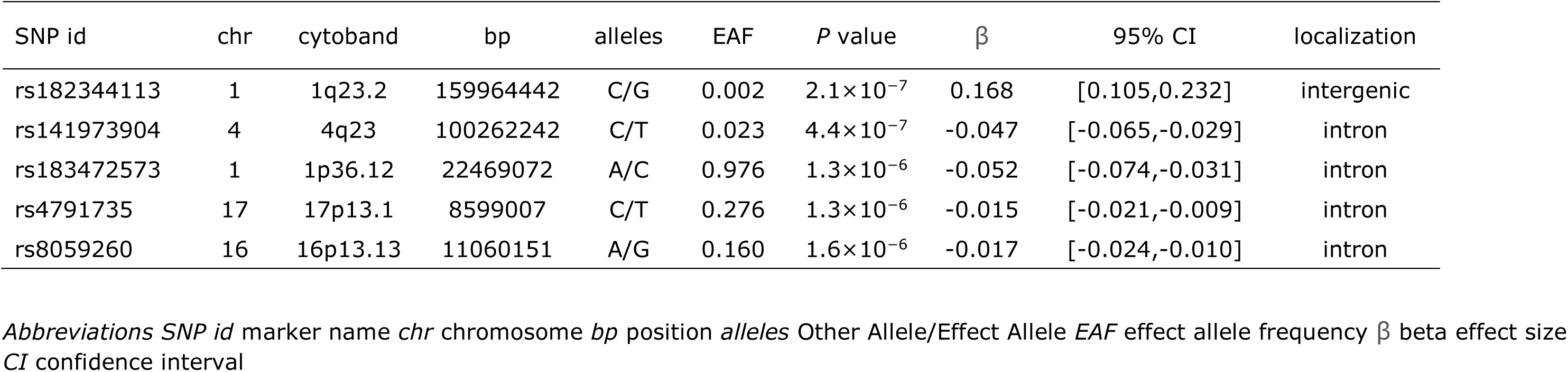
Common genetic variants for strongest associations (*P* < 10^−7^) with AUDIT

**Supplementary Table 7.**
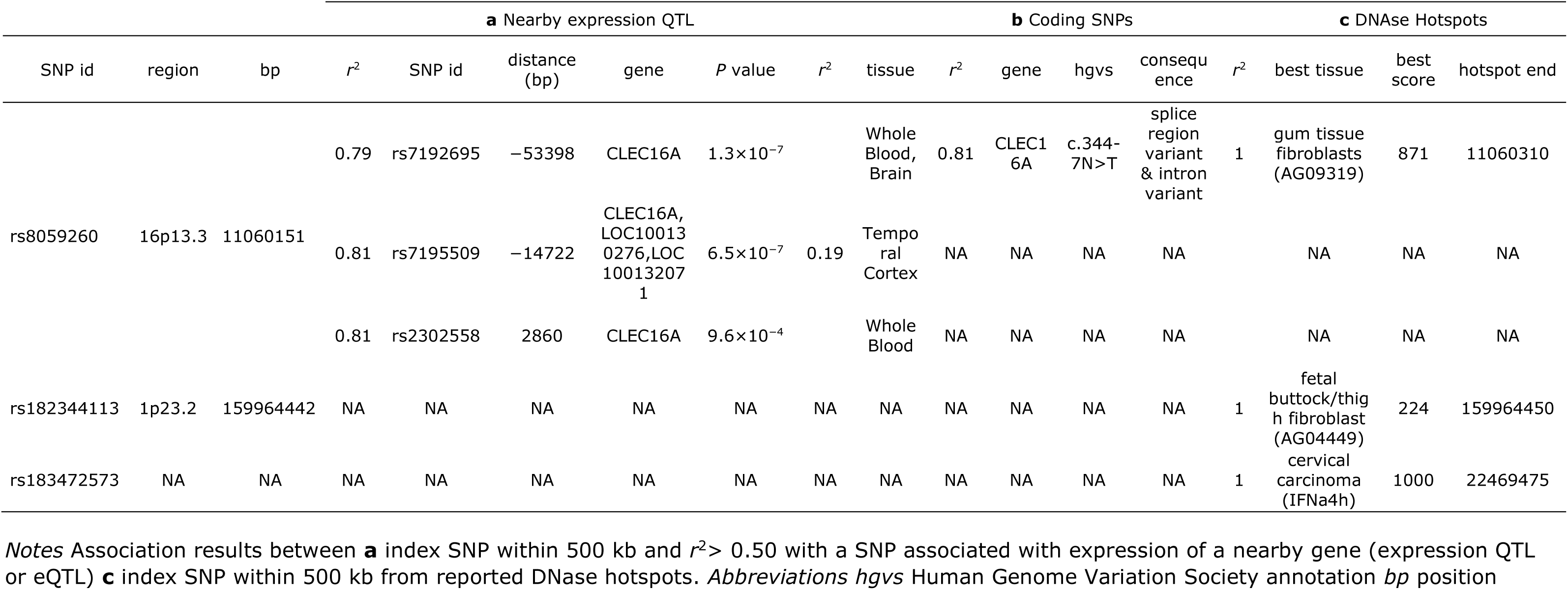
SNP-gene expression associations for the lead SNPs: **a** nerby expression QTL **b** nearby coding SNPs **c** nearby DNAse Hotspots

**Supplementary Table 8.** See Online Material for a comparison between previous single nucleotide polymorphisms associated with alcohol-related traits, as reviewed by (Bühler *et al*, 2015), and AUDIT scores

**Supplementary Table 9.** Extension of Figure 2 Genetic correlations between AUDIT scores and other relevant traits using LD Regression Score (LDSC)

| Phenotype | Sample Size | Reference | Genetic correlation ($r_g$) | Standard error | *P* value | FDR < 0.05 |
| --- | --- | --- | --- | --- | --- | --- |
| Years of education | 293,723 | (Okbay *et al*, 2016b) | 0.27 | 0.06 | 3.14E-05 | Yes |
| BMI | 339,224 | (Locke *et al*, 2015) | -0.25 | 0.06 | 1.48E-04 | Yes |
| ADHD | 53,293 | (Demontis *et al*, 2017) | -0.29 | 0.01 | 1.43E-03 | Yes |
| Smoking lifetime | 74,035 | (Furberg *et al*, 2010) | 0.42 | 0.13 | 1.52E-03 | Yes |
| Obesity class 1 | 98,697 | (Berndt *et al*, 2013) | -0.23 | 0.08 | 2.06E-03 | Yes |
| Alcohol consumption | 70,460 | (Schumann *et al*, 2016) | 0.68 | 0.23 | 3.40E-03 | Yes |
| Years of education | 101,069 | (Rietveld *et al*, 2013) | 0.27 | 0.09 | 6.08E-03 | Yes |
| Childhood IQ | 12,441 | (Benyamin *et al*, 2014) | 0.42 | 0.16 | 6.26E-03 | Yes |
| College attainment | 95,427 | (Rietveld *et al*, 2013) | 0.26 | 0.10 | 8.11E-03 | Yes |
| Cannabis lifetime use | 32,330 | (Stringer *et al*, 2016) | 0.38 | 0.17 | 2.60E-02 | No |
| Daily cigarettes | 68,028 | (Furberg *et al*, 2010) | -0.31 | 0.16 | 6.19E-02 | No |
| Alcohol dependence | 7,280 | (Gelernter *et al*, 2014) | 0.68 | 0.37 | 6.41E-02 | No |
| Intracranial brain volume | 13,171 | (Hibar *et al*, 2015) | 0.33 | 0.20 | 8.91E-02 | No |
| Anorexia nervosa | 17,767 | (Boraska *et al*, 2014) | 0.21 | 0.14 | 1.29E-01 | No |
| Loneliness | 10,760 | (Gao *et al*, 2017) | -0.39 | 0.31 | 2.08E-01 | No |
| Hippocampus | 30,717 | (Hibar *et al*, 2015) | -0.21 | 0.18 | 2.38E-01 | No |
| Childhood obesity | 13,848 | (Bradfield *et al*, 2012) | -0.12 | 0.12 | 3.04E-01 | No |
| Schizophrenia | 77,096 | (Ripke *et al*, 2014) | 0.05 | 0.07 | 4.95E-01 | No |
| Neuroticism | 170,911 | (Okbay *et al*, 2016a) | 0.06 | 0.09 | 5.08E-01 | No |
| Former smoker | 70,675 | (Furberg *et al*, 2010) | -0.11 | 0.18 | 5.58E-01 | No |
| Bipolar disorder | 16,731 | (Sklar *et al*, 2011) | 0.05 | 0.10 | 6.11E-01 | No |
| Extraversion | 63,030 | (van den Berg *et al*, 2016) | 0.07 | 0.16 | 6.43E-01 | No |
| Neuroticism | 106,716 | (Smith *et al*, 2016) | -0.04 | 0.09 | 6.57E-01 | No |
| Height | 253,288 | (Wood *et al*, 2014) | 0.02 | 0.05 | 7.12E-01 | No |
| Major depressive disorder | 18,759 | (Ripke *et al*, 2013) | 0.06 | 0.17 | 7.22E-01 | No |
| Depressive symptoms | 161,460 | (Okbay *et al*, 2016a) | -0.04 | 0.11 | 7.38E-01 | No |
| Nucleus accumbens | 30,717 | (Hibar *et al*, 2015) | 0.08 | 0.24 | 7.43E-01 | No |
| Caudate | 30,717 | (Hibar *et al*, 2015) | -0.02 | 0.13 | 8.87E-01 | No |
| Neo-openness to experience | 17,375 | (de Moor *et al*, 2012) | 0.02 | 0.18 | 8.93E-01 | No |
| Smoking age onset | 47,961 | (Furberg *et al*, 2010) | 0.01 | 0.22 | 9.40E-01 | No |
| Subjective wellbeing | 298,420 | (Okbay *et al*, 2016a) | -0.01 | 0.11 | 9.64E-01 | No |

## Extended Methods

A maximal set of unrelated individuals was chosen for the analysis using a segmental identity-by-descent (IBD) estimation algorithm (Henn *et al*, 2012) to ensure that only unrelated individuals were included in the sample. Individuals were defined as related if they shared more than 700 cM IBD, including regions where the two individuals shared either one or both genomic segments identical-by-descent. This level of relatedness (~20% of the genome) and corresponds approximately to the minimal expected sharing between first cousins in an outbred population.

We imputed participant genotype data against the September 2013 release of 1000 Genomes phase 1 version 3 reference haplotypes. We phased and imputed data for each genotyping platform separately. We phased using an internally developed phasing tool, Finch, which implements the Beagle haplotype graph-based phasing algorithm (Browning and Browning, 2007), modified to separate the haplotype graph construction and phasing steps. In preparation for imputation, we split phased chromosomes into segments of no more than 10,000 genotyped SNPs, with overlaps of 200 SNPs. We excluded SNPs with Hardy-Weinberg equilibrium *P*<10^−20^, call rate < 95%, or with large allele frequency discrepancies compared to European 1000 Genomes reference data. Frequency discrepancies were identified by computing a 2x2 table of allele counts for European 1000 Genomes samples and 2000 randomly sampled 23andMe research participants with European ancestry, and identifying SNPs with a □^2^ *P* < 10^−15^. We imputed each phased segment against all-ethnicity 1000 Genomes haplotypes (excluding monomorphic and singleton sites) using Minimac2 (Fuchsberger *et al*, 2015), using 5 rounds and 200 states for parameter estimation. After quality control, we analyzed 11,508,740 SNPs.

For the X chromosome, we built separate haplotype graphs for the non-pseudoautosomal region and each pseudoautosomal region, and these regions were phased separately. We then imputed males and females together using Minimac2, as with the autosomes, treating males as homozygous pseudo-diploids for the non-pseudoautosomal region.

For tests using imputed data, we use the imputed dosages rather than best-guess genotypes. We imputed HLA allele dosages from SNP genotype data using HIBAG (Zheng *et al*, 2014). We imputed alleles for HLAA, B, C, DPB1, DQA1, DQB1, and DRB1 loci at four-digit resolution. To test associations between HLA allele dosages and phenotypes, we performed linear regression using the same set of covariates used in the SNP based GWAS. We performed separate association tests for each imputed allele.

